# Antagonistic epistasis of Hnf4α and FoxO1 networks through enhancer interactions in β-cell function

**DOI:** 10.1101/2020.07.04.187864

**Authors:** Taiyi Kuo, Wen Du, Yasutaka Miyachi, Prasanna K. Dadi, David A. Jacobson, Domenico Accili

**Affiliations:** Department of Medicine and Berrie Diabetes Center, Columbia University College of Physicians and Surgeons, New York, NY; Department of Molecular Physiology and Biophysics, Vanderbilt University, Nashville, TN

## Abstract

Genetic and acquired abnormalities contribute to pancreatic β-cell failure in diabetes. Transcription factors Hnf4α (MODY1) and FoxO1 are respective examples of these two components, and are known to act through β-cell-specific enhancers. However, their relationship is unclear. Here we show by genome-wide interrogation of chromatin modifications that FoxO1 ablation in mature β-cells leads to increased selection of FoxO1 enhancers by Hnf4α. To model the functional significance we generated single and compound knockouts of FoxO1 and Hnf4α in β-cells. Single knockout of either gene impaired insulin secretion in mechanistically distinct fashions. Surprisingly, the defective β-cell secretory function of either single mutant in hyperglycemic clamps and isolated islets treated with various secretagogues, was completely reversed in double mutants. Gene expression analyses revealed the reversal of β-cell dysfunction with an antagonistic network regulating glycolysis, including β-cell “disallowed” genes; and that a synergistic network regulating protocadherins emerged as likely mediators of the functional restoration of insulin secretion. The findings provide evidence of antagonistic epistasis as a model of gene/environment interactions in the pathogenesis of β-cell dysfunction.

## Introduction

Genetic predisposition contributes to type 2 diabetes through changes in the expression and activity of transcription factors essential for β-cell function and insulin sensitivity (Ferrannini, 2010). The best characterized examples are the MODY gene mutations, which generally act in autosomal dominant fashion to impair insulin secretion (Fajans and Bell, 2011). In the vast majority of patients, however, such predisposition is more subtle, and requires acquired abnormalities, often associated with altered nutrient utilization, to give rise to overt β-cell dysfunction. An example of acquired transcriptional abnormalities leading to altered β-cell function is FoxO1, a nutrient-regulated transcription factor whose nutrient excess-driven failure leads first to metabolic inflexibility (Kim-Muller et al., 2014), and then to outright β-cell dedifferentiation (Cinti et al., 2016; Sun et al., 2019; Talchai et al., 2012). This is achieved in part by direct actions on glucose metabolism (Kuo et al., 2019b) and in part by regulating lineage stability (Kuo et al., 2019a).

We have been interested in exploring the functional relation between FoxO1 and Hnf4α, as a model of gene/environment interactions in the pathogenesis of β-cell dysfunction. When we ablated FoxO1, 3a and 4 in pancreatic progenitors or mature β-cells, gene ontology analyses revealed that Hnf4α targets were the most altered (Kim-Muller et al., 2014). However, whether these changes resulted in activation or repression of Hnf4α, and whether they contributed to, or offset FoxO1-induced β-cell dysfunction, was not clear.

Mutations of *HNF4A* cause MODY1 (Yamagata et al., 1996). Most patients develop diabetes in the third decade of life owing to a primary defect in insulin secretion (Herman et al., 1994) that can be treated long-term with sulfonylureas (Gardner and Tai, 2012). In mice, pancreas-wide (Pdx1-*Cre*) or mature β-cell (Rip-*Cre*) deletion of Hnf4α causes glucose intolerance due to defective insulin secretion (Boj et al., 2010; Gupta et al., 2005; Miura et al., 2006). There are shared phenotypic features between FoxO-deficient and Hnf4α-deficient β-cells (Gupta et al., 2005; Kim-Muller et al., 2014; Miura et al., 2006) that include compromised glucose-stimulated insulin secretion, and altered Pparα and Hnf1α gene network expression, indicative of shared pathways and targets of both transcription factors.

The present studies were undertaken to analyze a potential FoxO1/Hnf4α epistasis. Surveys of histone H3 lysine 27 acetylation for active promoters and enhancers uncovered an interplay between Hnf4α and FoxO1 at β-cell enhancers. Thus, we generated and compared mutant animals lacking FoxO1, Hnf4α, or both in β-cells. Surprisingly, FoxO1/Hnf4α double knockout mice (DKO) showed restored glucose tolerance and insulin secretion compared to single knockouts. These findings fundamentally change our understanding of the interactions within transcriptional networks regulating β-cell function by highlighting a heretofore unrecognized antagonistic epistasis (Snitkin and Segre, 2011).

## Results

### Altered enhancer selection in β-cell failure

Activation of FoxO1 enhancers is a key event in β-cells in response to insulin-resistant diabetes in *db/db* mice (Kuo et al., 2019b). We investigated FoxO1-regulated enhancers in a multiparity model of diabetes associated with genetic ablation of FoxO1 (Kuo et al., 2019a; Talchai et al., 2012). To this end, we subjected fluorescently sorted β-cells from multiparous diabetic β-cell-specific FoxO1 knockout mice (DM) and glucose-tolerant controls (NGT) to genome-wide chromatin immunoprecipitation with acetylated histone 3 lysine 27 antibodies (H3K27ac) as a marker of active enhancers and promoters (Creyghton et al., 2010). DM mice show evidence of multiparity-associated β-cell failure with random as well as re-fed hyperglycemia as well as fasting and refed hypoinsulinemia (Kuo et al., 2019a; Talchai et al., 2012). We observed a global *increase* of H3K27ac in active regions (Fig. S1A), and in individual genes (Fig. S1B) of DM β-cells. Nearly half (44%) of H3K27ac peaks were located in introns, 14% in intergenic regions, and 16% in promoters (Fig. S1C). We analyzed the data based on H3K27ac regions associated with promoters (defined as +/– 3kb from transcription start sites) or distal enhancers (defined as beyond promoters) (Lenhard et al., 2012). Interestingly, we detected significant changes in distal enhancers in the absence of FoxO1, regardless of age or parity (Fig. 1A). In contrast, we found minimal alterations in the promoters (Fig. 1B). These data confirm the importance of FoxO1 in distal enhancers (Kuo et al., 2019b).

**Figure 1.**
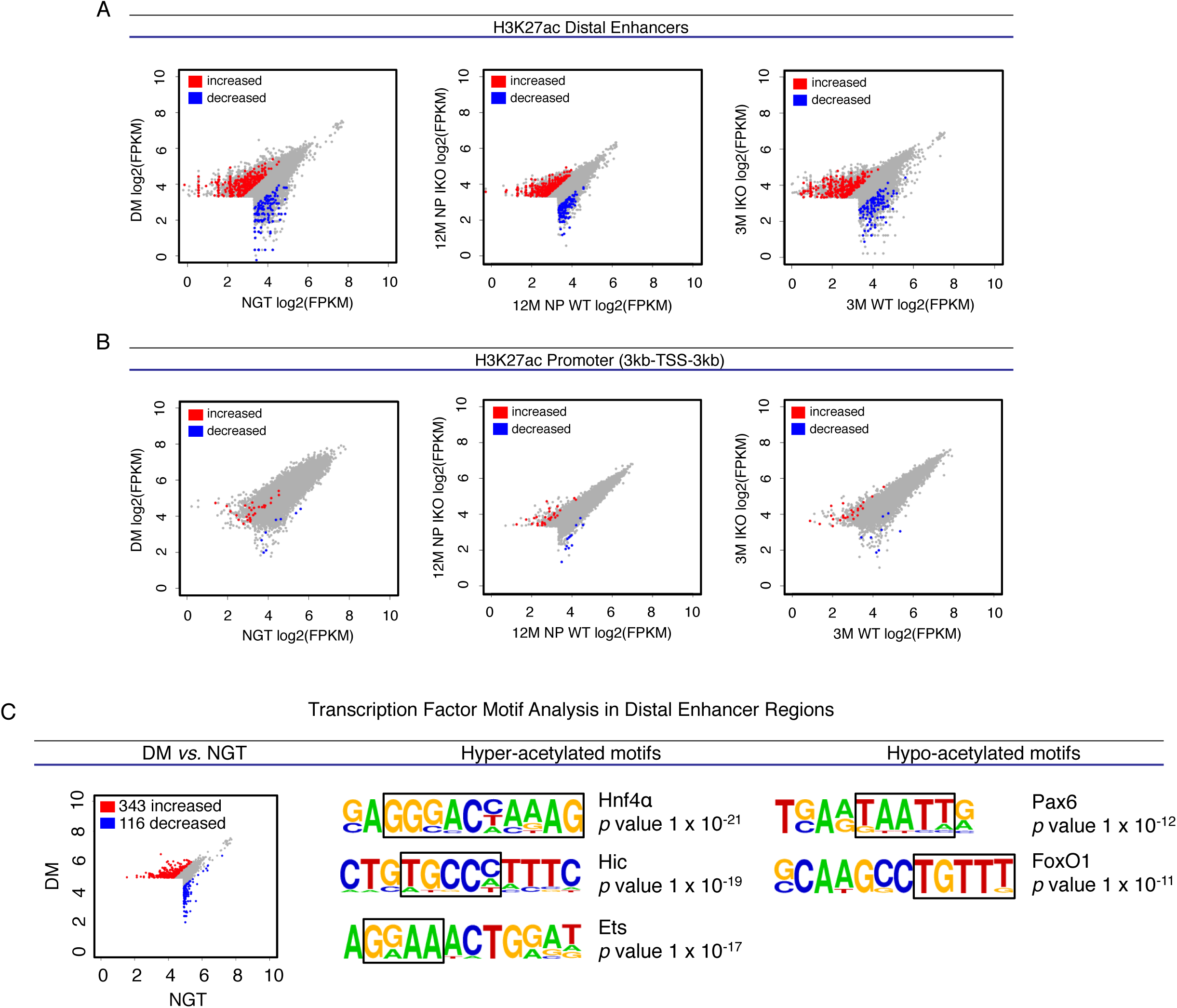
Motif analysis of distal enhancers. **(A-B)** Scatterplots of H3K27ac regions in distal enhancers (A) and promoters (B) in WT and IKO β-cells isolated under three conditions: diabetic multiparous IKO (DM) and WT controls (normal glucose tolerance, or NGT); age-matched nulliparous (12M NP) IKO and WT; and 3-month-old (3M) IKO and WT controls. There are 417 up-regulated (red) and 149 down-regulated (blue) H3K27ac regions in distal enhancers (A) and 31 up-regulated and 15 down-regulated H3K27ac regions in promoters (B) shared among the three comparisons. **(C)** Transcription factor motif analysis in differentially acetylated distal enhancers between DM and NGT. For each transcription factor motif, consensus or partial consensus sequence is boxed, and *p* value is shown.

To investigate the functional consequences of the altered distal enhancer profile of DM β-cells, we mined transcription factor binding motifs in regions with increased or decreased H3K27ac marks in DM *vs*. NGT. We reckoned that activation of transcription factors would be associated with enrichment of the relevant motifs in hyper-acetylated enhancers, whereas suppression of transcription factors would be associated with enrichment of their motifs in hypo-acetylated enhancers (Kuo et al., 2019b). FoxO1 was the second most-underrepresented motif found within hypo-acetylated enhancers in DM β-cells, thus validating the analysis (Fig. 1C). We also found a significant depletion of the Pax6 motif, a FoxO1 target (Kim-Muller et al., 2016b; Kitamura et al., 2009) that regulates *Ins1, Ins2* (Sander et al., 1997), Nkx6.1, Pdx1, Slc2a2 and Pc1/3 in β-cells (Hart et al., 2013). Pax6 represses alternative islet cell genes including ghrelin, glucagon, and somatostatin (Swisa et al., 2017), and ablation of Pax6 in the adult pancreas leads to glucose intolerance due to β-cell dedifferentiation (Hart et al., 2013; Swisa et al., 2017). The data are consistent with the possibility that combined abnormalities of FoxO1 and Pax6 contribute to β-cell dedifferentiation.

In contrast, Hnf4α was the top hyper-acetylated motif with increased H3K27ac in DM β-cells (Fig. 1C), consistent with prior gene ontology analyses indicating activation of the Hnf4α network in triple FoxO (Kim-Muller et al., 2014) and single FoxO1 knockout β-cells (Kuo et al., 2019a). In addition, Hic (Liu et al., 2011) and Ets (Luo et al., 2014), two networks of potential interest for β-cell stability, were over-represented. Interestingly, Ets1 overexpression in β-cells suppresses insulin secretion (Chen et al., 2016). These data reveal a striking rearrangement of active enhancers in diabetic β-cells.

### Restoration of metabolic parameters in FoxO1 and Hnf4α double knockout mice

To examine whether the increased representation of Hnf4α enhancers is compensatory or contributory vis-à-vis β-cell dysfunction, we generated β-cell-specific FoxO1 (IKO), Hnf4α (HKO) (Miura et al., 2006) as well as double knockout (DKO) mice using Rip-*Cre* transgenic mice (Herrera, 2000) that do not give rise to extra-pancreatic recombination or produce Gh mRNA (Brouwers et al., 2014). The animals were born to term in the expected Mendelian ratios, and showed no growth abnormalities (Fig. S2A).

We performed oral glucose tolerance tests. As previously reported, IKO or HKO mice displayed glucose intolerance (Fig. 2A) (Kuo et al., 2019a; Miura et al., 2006; Talchai et al., 2012). Unexpectedly, compared to single knockouts, DKO mice showed restoration of normal glucose tolerance (Fig. 2A and 2B). We also assessed plasma glucose levels within 5 minutes of intraperitoneal glucose injection as a surrogate measure of insulin secretion. Consistent with glucose tolerance tests, IKO mice showed elevated glucose levels within 5 minutes, whereas DKO mice showed no difference compared to WT (Fig. 2C). To investigate the nutrient response, we measured glucose and insulin levels in fasted and refed mice. While no difference was found in fasted plasma glucose and insulin levels between all 4 genotypes, refeeding led to elevated plasma glucose and reduced insulin levels in IKO mice compared to WT (Fig. 2D, 2E). Interestingly, DKO and WT mice show similar levels of glucose and insulin in response to a meal (Fig. 2D, 2E).

**Figure 2.**
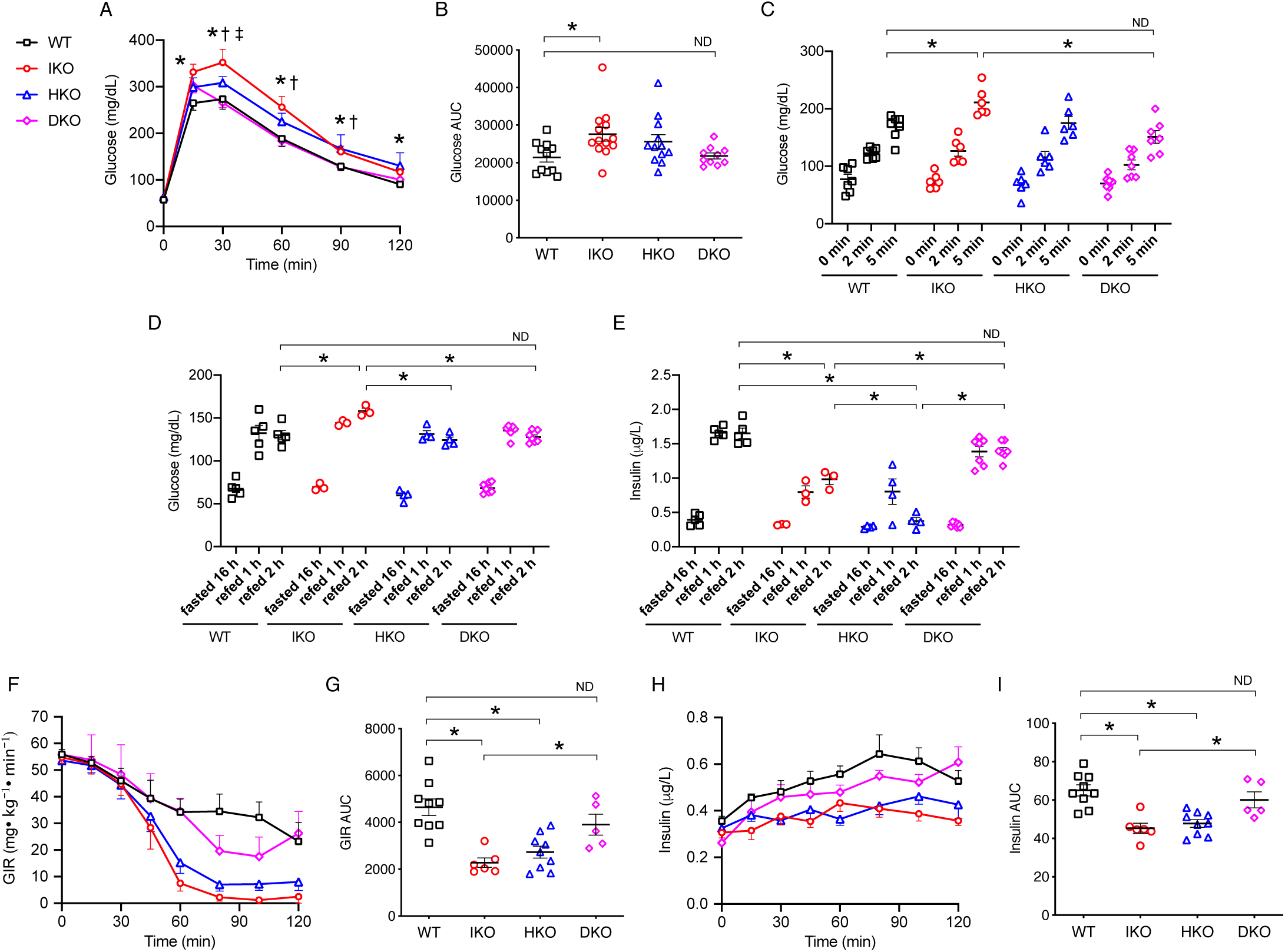
Glucose tolerance tests, fasting and refeeding response, and hyperglycemic clamps. **(A)** Intraperitoneal glucose tolerance test in WT (n=11), IKO (n=13), HKO (n=12), and DKO (n=10) mice. † represents *p* < 0.05 for IKO *vs.* DKO, * indicates *p* < 0.05 for WT *vs.* IKO, and ‡ shows *p* < 0.05 for HKO *vs.* DKO. **(B)** Area under the curve (AUC) in **A**. Error bars represent SEM, \**p* < 0.05 by Student’s *t* test and ANOVA. **(C)** Glucose levels in WT (n=7), IKO (n=6), HKO (n=5), and DKO (n=7) mice within 5 min of intraperitoneal glucose injection. **(D-E)** Glucose (D) and plasma insulin levels (E) in fasted or refed WT (n=5), IKO (n=3), HKO (n=4), and DKO (n=5) mice. **(F)** Glucose infusion rates (GIRs) during hyperglycemic clamps in WT (n=9), IKO (n=6), HKO (n=9), and DKO (n=5) mice, and **(G)** quantification of AUC. Error bars represent SEM, \**p* < 0.05 by Student’s *t* test and ANOVA. **(H)** Circulating insulin levels during hyperglycemic clamps in WT (n=9), IKO (n=6), HKO (n=9), and DKO (n=5) mice, and **(I)** quantification of AUC. Error bars represent SEM, \**p* < 0.05 by Student’s *t* test and ANOVA. ND stands for no difference.

Next, to assess insulin secretory capacity *in vivo*, we performed hyperglycemic clamps. By intravenous glucose administration, we raised glycemia to ∼350 mg/dL (Fig. S2B and S2C), and measured the rate of glucose infusion necessary to maintain this level. DKO showed increased glucose infusion rates compared to IKO and HKO, to a level indistinguishable from WT (Fig. 2F and 2G). Plasma insulin levels increased accordingly, consistent with an improved insulin secretory capacity in DKO mice (Fig. 2H and 2I). In contrast, all 4 genotypes responded to exogenous insulin injection to a comparable extent (Fig. S2D). These data are consistent with the possibility that FoxO1 and Hnf4α have antagonistic epistasis in β-cells, and that increased Hnf4α enhancer occupancy contributes to β-cell dysfunction in the absence of FoxO1.

### Secretagogue-stimulated insulin secretion and calcium flux

To investigate the mechanism of restored insulin secretion, we subjected purified islets from all four genotypes to insulin secretion in response to various secretagogues. Consistent with the *in vivo* studies, IKO and HKO islets showed impaired insulin secretion in response to high glucose or arginine, which was restored to WT levels in DKO islets (Fig. 3A and 3B). Similar to studies in triple FoxO knockout islets, IKO islets also showed compromised insulin secretion in response to the sulfonylurea glibenclamide and the PKC activator PMA (Kim-Muller et al., 2014). Both responses were normal in HKO, and were restored in DKO islets (Fig. 3C and 3D). Although the sulfonylurea response differs from previously reported data in Hnf4α knockout β-cells (Gupta et al., 2005; Miura et al., 2006), it is indeed consistent with the clinical response of MODY1 patients (Gardner and Tai, 2012).

**Figure 3.**
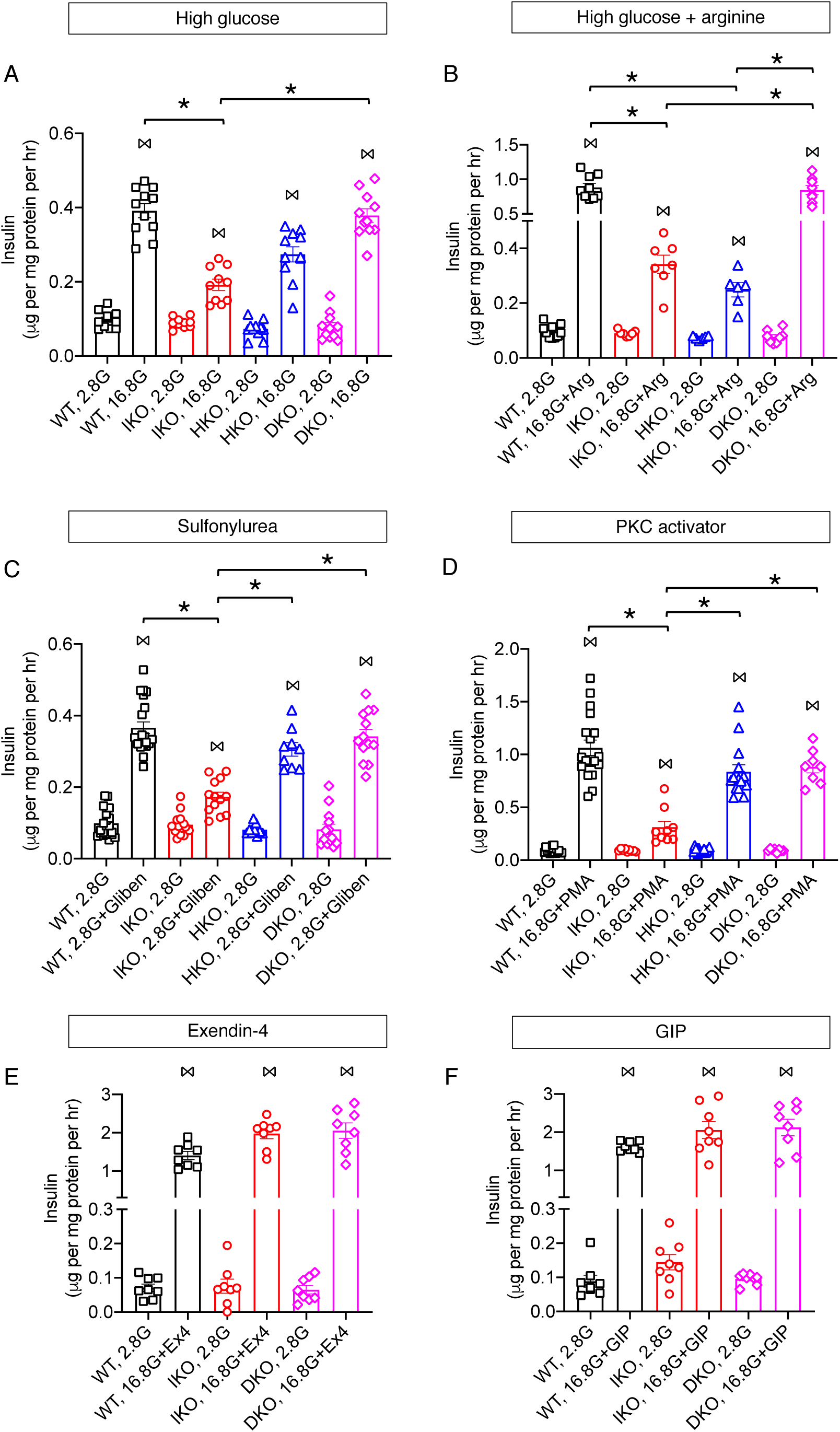
*Ex vivo* insulin secretion in purified islets. **(A)** High glucose- (16.8 mM, or 16.8G) stimulated insulin secretion in WT (n=11), IKO (n=10), HKO (n=11), and DKO (n=11) islets. **(B)** 16.8G and L-arginine- (10 mM, or Arg) stimulated insulin secretion in WT (n=10), IKO (n=7), HKO (n=6), and DKO (n=8) islets. **(C)** Glibenclamide- (1 μM) stimulated insulin secretion in WT (n=21), IKO (n=14), HKO (n=9), and DKO (n=13) islets. **(D)** PMA- (1 μM) stimulated insulin secretion in WT (n=18), IKO (n=9), HKO (n=14), and DKO (n=8) islets. **(E)** Exendin-4- (Ex4, 1 μM) stimulated insulin secretion in WT (n=8), IKO (n=8), and DKO (n=8) islets. **(F)** GIP- (1 μM) stimulated insulin secretion in WT (n=8), IKO (n=8), and DKO (n=8) islets. Islets were first incubated in basal glucose (2.8 mM, or 2.8G), followed by the indicated secretagogue(s). Results are normalized to total protein contents and duration of treatments. Error bars indicate SEM, ⋈ represents *p* < 0.05 when comparing basal glucose- and secretagogue-stimulated insulin secretion in the same genotype by Student’s *t* test, while * shows *p* < 0.05 when comparing secretagogue-stimulated insulin secretion in different genotypes by Student’s *t* test and ANOVA.

To probe the amplifying signal in insulin secretion, we tested exendin-4, a GLP-1 agonist, and GIP treatment. We found that both exendin-4- and GIP-stimulated insulin secretion were comparable in WT, IKO, and DKO islets (Fig. 3E and 3F), indicating that the GLP-1 and GIP receptor-dependent pathways of insulin secretion were not affected by FoxO1 ablation. Furthermore, it has been shown that GLP-1 can stimulate insulin secretion in HKO islets, although the magnitude was trending lower compared to WT (Miura et al., 2006).

Glycolysis-generated ATP binds to and closes K_ATP_ channels resulting in membrane depolarization and activation of voltage-dependent calcium channels (VDCCs). Calcium influx through VDCCs induces fusion of insulin granules to the β-cell plasma membrane and insulin exocytosis. We investigated calcium flux in islets of various genotypes. Unlike triple FoxO knockout islets (Kim-Muller et al., 2014), membrane depolarization-induced calcium influx in response to glucose and KCl was slightly increased in IKO (Fig. 4A, 4B). In addition to the genotype, differences in experimental design and time course of measures may account for this difference. Consistent with prior observations, HKO islets showed decreased magnitude and a delayed onset of calcium influx when compared to WT islets (Fig. 4A, 4B). Interestingly, DKO restored the magnitude but not the delayed onset of calcium flux. As a result, the area under the curve of the calcium responses following glucose-stimulation was restored by DKO at later, but not earlier time points (Fig. 4A, 4B). In sum, FoxO1 and Hnf4α affect insulin secretion in mechanistically distinct ways and result in antagonistic epistatic interactions.

**Figure 4.**
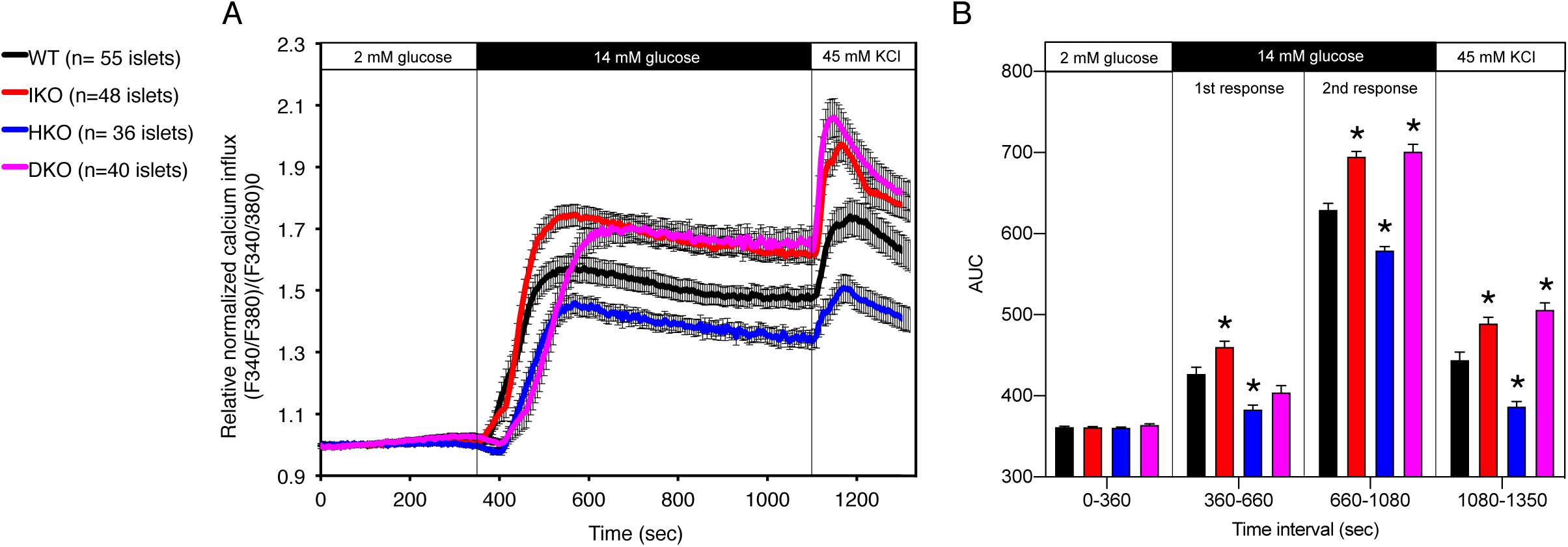
Average whole islet calcium response. Islets were incubated for 20 minutes in 2 mM glucose-containing solution and imaged using FURA2 prior to treatment with 14 mM glucose. **(A)** Representative Fura-2 AM recordings (F340/F380) of changes in whole-islet intracellular calcium flux for WT, IKO, HKO, and DKO mice. The secretagogues employed are indicated on top. **(B)** Area under the curve (AUC) for different secretagogue-stimulated insulin secretion in A. Error bars indicate SEM, * represents *p* <0.05 when comparing to WT by Student’s *t* test and ANOVA.

### Different mechanisms of β-cell dysfunction in FoxO1 *vs.* Hnf4α mutants

To understand the mechanisms of epistasis between FoxO1 and Hnf4α, we introduced a fluorescent *ROSA26*-Tomato reporter in the various mutant strains and used it to sort genetically-labeled Tomato-positive β-cells (Kuo et al., 2019a). We performed RNAseq in WT, IKO, HKO, and DKO β-cells and identified expression changes by differential gene-calling. Our working hypothesis was that functional antagonistic epistasis could result from: (i) synergistic regulation of networks that inhibit insulin secretion; (ii) antagonistic regulation of networks that promote insulin secretion; (iii) mixed regulation of distinct components of individual networks (Fig. 5) (Snitkin and Segre, 2011).

**Figure 5.**
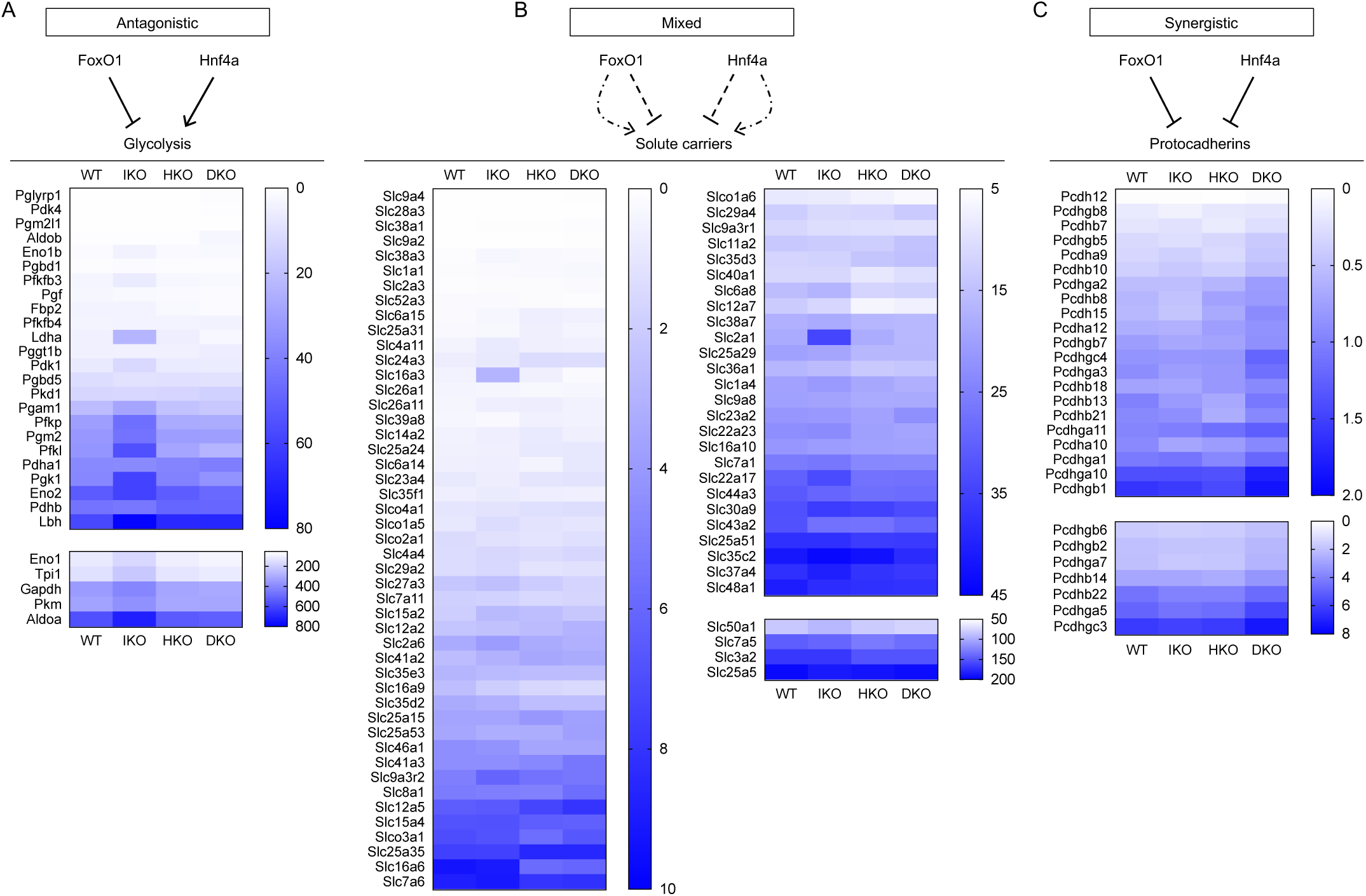
Epistatic modalities between FoxO1 and Hnf4α. **(A)** Antagonistic epistasis-regulated glycolytic network. **(B)** Mixed epistasis for solute carrier lineage. **(C)** Synergistic epistasis-controlled protocadherins. Heatmaps demonstrate altered gene expression in DKO *vs*. IKO or HKO β-cells. Scale bars indicate FPKM.

When compared to WT, there were changes to 769 mRNAs in IKO β-cells (Table S1), 1,327 in HKO (Table S2), and 1,692 in DKO (Table S3). Thus, the restoration of insulin secretion in DKO seemingly is not due to a normalization of gene expression, but rather to the emergence of a distinct gene expression profile. This is confirmed by comparisons of single with double mutants, where DKO β-cells displayed 1,787 mRNA changes compared to IKO (Table S4), and 666 compared to HKO (Table S5). Furthermore, there were 1,165 mRNA changes between IKO and HKO β-cells (Table S6).

In IKO β-cells, we detected the expected alterations of known FoxO1 target mRNAs, with increased glycolytic genes (Buteau et al., 2007), including β-cell “disallowed” genes (Kuo et al., 2019b) and *Aldh1a3* (Kim-Muller et al., 2016a), as well as decreased *Cyb5r3* (Fan et al., 2020) (Table S1). These data are consistent with previous findings indicating that triple FoxO ablation results in altered metabolic coupling (Kim-Muller et al., 2014). Specifically, the combination of increased glycolysis and impaired coupling to oxidative phosphorylation by activation of disallowed genes may result in lower efficiency of ATP production and increased ROS generation (Robertson et al., 2003).

Previous reports differ with regard to gene expression patterns associated with Hnf4α deletion in β-cells (Gupta et al., 2005; Miura et al., 2006). In our cross, the most striking gene expression changes in HKO were the substantial increases of abundant mRNAs encoding mitochondrial cytochrome components (COX, CYTB, ND1, ND2, ND4, and ND5) (Table S2). Conversely, the mitochondrial complex III oxidoreductase *Cyb5r3* was decreased, possibly resulting in altered electron transport across the mitochondrial chain (Fan et al., 2020). There were extensive increases in the family of KCN-type potassium channels, which may affect membrane depolarization and insulin secretion, and thus explain the preserved response to sulfonylurea (Fig. S3A). The levels of L-type VDCC *Cacnb3* mRNA were decreased, a finding that may partly explain the impaired calcium response in HKO islets (Table S2). Interestingly, this change was not reversed in DKO, consistent with the observation that first-phase calcium flux is NOT restored, despite the overall improvement of calcium flux and insulin secretion (Table S3). These data indicate that the underlying mechanisms of islet dysfunction differ in IKO *vs*. HKO islets. There were two notable exceptions to this: *Cyb5r3*, as indicated above, and *Gpr119*, an orphan GPCR that is mechanistically linked to insulin secretion (Abdel-Magid, 2019), were equally decreased in both IKO and HKO (Table S1 and S2).

### Epistatic network analysis

Clues as to the potential mechanism of reversal of β-cell dysfunction emerged from comparing single mutants with DKO as well as WT (Table S3). The glycolytic pathway, including key β-cell “disallowed” genes, such as *Ldha* and *Slc16a3*, was substantially decreased in DKO and WT compared to IKO. Indeed, DKO levels were even lower than WT islets. The predicted outcome of this change would be a decrease in glucose utilization and tighter coupling of glycolysis with oxidative phosphorylation, owing to reduced *Ldha* activity and monocarboxylate transport. This change is consistent with synergistic antagonism of FoxO1 and Hnf4α on the glycolytic cascade, with the former acting as a suppressor and the latter as an activator of this pathway (Fig. 5A, 5B).

In contrast, changes to HKO gene expression patterns were not reversed in either WT or DKO islets. For example, the patterns of KCN expression remained virtually identical (Fig. S3A). Even when analyzing the vast and heterogenous family of solute carriers (*Slc*), 61% of the changes seen in HKO failed to be reversed, as opposed to 23% in IKO islets. Thus, the majority of Hnf4α targets are not epistatic with FoxO1. However, a remarkable synergistic regulation of the protocadherin gene family emerged by comparing HKO with WT or DKO islets (Fig. 5C). Protocadherins are thought to participate in the formation of adherens (tight) junctions between β-cells that facilitate calcium influx and fusion of secretory vesicles (Dissanayake et al., 2018). This may explain the unique patterns of calcium entry in DKO islets, which are slower but reach higher levels than WT. This regulation of the protocadherin family is consistent with a model of synergistic epistasis. In this instance, both FoxO1 and Hnf4α act in a similar fashion to repress expression of these genes, and it’s only when both are removed that the full extent of the regulation becomes apparent.

A second example of synergistic epistasis between FoxO1 and Hnf4α were mRNAs encoding non-β-cell hormones (Fig. S3B). *Gcg, Sst, Npy, Ppy*, and *Pyy* increased substantially in DKO islets compared to IKO, consistent with reduced lineage stability, a feature of “resting” or dedifferentiated β-cells (Buteau et al., 2007; Talchai et al., 2012; Talchai and Accili, 2015).

### Relevance to human type 2 diabetes GWAS

To investigate the human disease relevance of our findings, we compared DKO with either single IKO mutants or WT. We integrated enhancer occupancy and RNAseq data to select genes with increased H3K27ac marks and mRNA levels, or decreased H3K27ac and mRNA levels. We selected genes concordantly up- or down-regulated in DKO compared to both WT *and* IKO as potential “compensatory” genes, and queried a database of genetic variations in human type 2 diabetes GWASdb SNP (Rouillard et al., 2016) (URL: http://amp.pharm.mssm.edu/Harmonizome) to identify those associated with type 2 diabetes (Tables S7 and S8).

Genes with increased mRNA and H3K27ac numbered 12 in the DKO vs. WT, and 7 in the DKO vs. IKO comparison. Two (*Svil* and *Prex1*) were shared in common. Genes with decreased mRNA and H3K27ac were 22 in DKO vs. WT, and 32 in DKO vs. IKO β-cells (Tables S7 and S8, Fig. S4). The latter included *ARHGEF9*, one of only 20 genes concordantly decreased in three independent RNA profiling datasets of type 2 diabetes patients (Zhong et al., 2019). 12 genes (*Ttc22, Sel1l3, Rspo4, Pla2g4b, Nr1h4, Lad1, Jmjd7, Fut1, Elovl2, Dll4, Cacnb3*, and *Baiap2l2*) were shared in common between the two comparisons (Table S7 and S8). Altogether, three genes were associated with human variant sequences conferring an increased risk of type 2 diabetes: *Svil, Prex1*, and *Elovl2*.

## Discussion

β-cell function is maintained through a restricted network of transcription factors (Fujitani, 2017). However, the redundancy, heterogeneity, and complexity of gene expression patterns associated with experimental or naturally occurring perturbations in the activity of these “master” regulators complicate the task of interrogating the mechanisms of dysfunction resulting from their interactions. The main findings of this work are: i) FoxO1 deletion results in a redistribution of Hnf4α to FoxO1 distal enhancers in β-cells; ii) this redistribution underlies an antagonistic epistasis between these transcription factors affecting insulin secretion; iii) gene expression analyses reveal different epistatic modalities regulating different networks, examples of which include antagonistic epistasis of the glycolysis pathway, and synergistic epistasis of the protocadherin gene family. Epistatic synergism between paralogues in β-cell function has been proposed (Boj et al., 2010). But antagonistic epistasis between distinct factors through cis-acting enhancers and resulting in mechanistically distinct phenotypes had not been disclosed.

The interaction between FoxO1 and Hnf4α occurs at multiple levels. The two proteins physically interact in liver (Puigserver et al., 2003). In addition, Hnf4α-initiated glucokinase transcription is repressed by FoxO1, acting as a co-repressor (Ganjam et al., 2009; Hirota et al., 2008) by way of SIN3a (Langlet et al., 2017). However, the co-regulation of pancreatic β-cell distal enhancers by FoxO1 and Hnf4α has not been revealed until now, and its relevance to the progression of common forms of diabetes in humans will have to be further explored.

Type 2 diabetes is a polygenic disease, and β-cell failure has acquired and genetic components. FoxO1 can be considered an example of the former, and Hnf4α of the latter. Both homozygous null mutations in β-cells are associated with an insulin secretory defect, although the mechanisms appear to differ. Thus, the absent sulfonylurea response in FoxO1 knockout mice is reminiscent of the acquired failure seen in diabetic patients, while Hnf4α knockout show a preserved sulfonylurea response, like MODY1 patients (Gardner and Tai, 2012). Likewise, the calcium response to glucose differs in the two mutants, although this may partly be explained by the compensatory function of FoxO3 in single FoxO1 knockouts (Kim-Muller et al., 2014).

Evidence of antagonistic epistasis between these factors arises from the surprising phenotype of the double mutant mice. However, the majority of Hnf4α-dependent changes are unaffected by the FoxO1 mutation, indicating that epistasis between these two proteins occurs primarily within the FoxO1 regulome. For example, the Hnf1α network is inhibited in HKO mutants (data not shown), and this alteration is not rescued in DKO mice. Conversely, most changes induced by FoxO1 ablation appear to be offset by the Hnf4α mutation. The most striking example is the antagonistic regulation of glycolysis, where FoxO1 acts as a suppressor and Hnf4α as an activator of gene expression. Although the association of reduced expression of glycolytic genes with restored insulin secretion in DKO islets may appear counterintuitive, it includes suppression of *Lhda*, and therefore can be conducive to better metabolic coupling of glycolysis with mitochondrial oxidative phosphorylation, leading to β-cell “rest” and reduced ROS production (Robertson et al., 2003).

A second example of epistatic network is represented by the synergistic suppression of protocadherins. The release of this suppression in DKO mice may lead to tighter coupling of neighboring β-cells and therefore to faster propagation of action potentials, reducing heterogeneity among β-cells (Dissanayake et al., 2018). Consistent with this hypothesis is the observation that calcium influx into DKO occurs even more rapidly than in WT β-cells. While further studies will be required to investigate this mechanism, these observations raise the possibility that agents that improve cell-cell communication can restore insulin secretion in diabetes.

An additional mechanism connecting the functional restoration of insulin secretion to changes in intracellular vesicle trafficking emerges from the integration of RNAseq with enhancer occupancy and human diabetes GWAS or RNAseq data. We found alterations of *Arhgef9*, encoding Rho guanine nucleotide exchange factor 9, a guanine exchange factor for Cdc42. This is one of the few genes consistently identified in multiple surveys of diabetic islet gene expression (Zhong et al., 2019). Small G-proteins, such as Rho, Rac1, Cdc42, and Arf6 play important roles in cytoskeletal remodeling, vesicle fusion, and insulin exocytosis (Kowluru, 2017). They cycle between inactive (GDP-bound) and active (GTP-bound) conformations, regulated by a combination of activating guanine nucleotide exchange factors (GEFs), inactivating GTPase-activating proteins (GAPs), and inhibitory GDP-dissociation inhibitors (GDI). In particular, Cdc42 depletion leads to a loss of second-phase insulin secretion, while Cdc42 activation is selectively responsive to glucose but not KCl (Wang et al., 2007). Thus, Arhgef9/Cdc42-dependent cytoskeletal rearrangement can be a mechanism to rescue the secretory phenotype in DKO *vs.* IKO β-cells.

Two of the genes associated with diabetes susceptibility in human GWAS studies and dysregulated in IKO or DKO islets are also related to cytoskeletal remodeling. *Prex1* (phosphatidylinositol-3,4,5-triphosphate-dependent Rac exchange factor 1) activates Rac1, leading to cytoskeletal rearrangement (Balamatsias et al., 2011). β-cell-specific Rac1 knockout mice display reduced glucose-stimulated insulin secretion due to failure to depolymerize F-actin, which prevents recruitment of insulin granules and insulin secretion (Asahara et al., 2013). *Svil* (encoding Supervillin) bridges membranes with the actin cytoskeleton (Chen et al., 2003). The combined increases in Arhgef9-Cdc42 and Prex1-Rac1 provide a potential link between the Rho GTPase family and cytoskeletal remodeling-mediated insulin secretion, and thus a compensatory mechanism to explain the pattern of calcium influx in DKO β-cells. F-actin forms a web beneath the plasma membrane that is often viewed as a barrier to the release of insulin granules in basal conditions (Kalwat and Thurmond, 2013; Orci et al., 1972). The depolymerization of F-actin is essential for nutrient-stimulated, second phase insulin secretion, which requires recruitment of insulin granules from the intracellular storage pools to the plasma membrane. The cyclic nature of F-actin remodeling, regulated by Cdc42 and Rac1, is necessary for the regulation of insulin granule exocytosis (Kalwat and Thurmond, 2013).

In summary, increased occupancy of Hnf4α through its re-distribution to FoxO1 distal enhancers is contributory to β-cell dysfunction, highlighting the role of antagonistic epistasis between transcription factors in the maintenance of pancreatic β-cells. These findings can provide a model to understand the pathophysiology of β-cell dysfunction.

## Methods

### Animals

We obtained Hnf4α flox/flox mice from Dr. Frank J. Gonzalez (Miura et al., 2006), and crossed Rip-*Cre*/*Rosa26-Tomato*/floxed *FoxO1* mice with floxed *Hnf4α* (Kuo et al., 2019a) to generate β-cell-specific FoxO1 and Hnf4α knockout mice. We categorize mice with at least one allele of FoxO1 and Hnf4α as WT for DKO studies. All mice were fed chow diet, and maintained on a 12-h light cycle (lights on at 7am). Epigenetic screening was performed in female mice, and FoxO1 and Hnf4α double knockout validations were carried in male mice.

### Metabolic Parameters

We performed intraperitoneal glucose tolerance tests with glucose (2g/kg) after an overnight (16 h) fast, and insulin tolerance tests by injecting insulin (0.75 units/kg) after a 5-h fast (Kuo et al., 2016). For hyperglycemic clamps, we followed a previously described protocol (Kuo et al., 2019b). To assess insulin capacity *in vivo*, we raised the plasma glucose level to ∼350 mg/dL by continuous infusion of glucose, and maintained by adjusting the glucose infusion rate on the pump. Throughout the 120 min period, we recorded glucose infusion rates and plasma glucose levels, and collected plasma for insulin measurement.

### Fluorescence-assisted β-cell sorting

Pancreatic islets were isolated as described and allowed to recover in RPMI supplemented with 15% FBS overnight (Kuo et al., 2016). The next day, islets were washed with PBS, and dissociated with trypsin, followed by quenching with FBS. Thereafter, the dispersed islets were spun down and washed with PBS, and incubated with sytox red (Thermo Fisher, S34859) to identify live cells and DNase I (Sigma, D4513) on ice until sorting. Prior to sorting, cells were filtered through a 35-μm cell strainer (Corning 352235) to remove clusters. Tomato-positive and -negative cells differed by ∼2 orders of magnitude in tomato fluorescence with excitation of 554nm and emission of 581nm by Influx™ cell sorter (BD Biosciences).

### H3K27ac ChIPseq

Islet β-cells were genetically labeled with ROSA26-Tomato fluorescence with Rip-*Cre* allele, and FAC-sorted β-cells were used for histone H3K27ac ChIPseq with anti-H3K27ac antibody (Active Motif, 39133). ChIP and ChIPseq were performed as previously described (Kuo et al., 2019a; Kuo et al., 2012). The resulting DNA libraries were quantified with Bioanalyzer (Agilent), and sequenced on Illumina NextSeq 500 with 75-nt reads and single end. Reads were aligned to mouse genome mm10 using the Burrows-Wheeler Aligner (BWA) algorithm with default settings (Li and Durbin, 2009). These reads passed Illumina’s purity filter, and aligned with no more than 2 mismatches. Duplicate reads were removed, and only uniquely mapped reads with a mapping quality greater or equal to 25 were subjected for further analysis. Alignments were extended in silico at their 3’ ends to a length of 200-bp, which is the average genomic fragment length in the size-selected library, and assigned to 32-nt bins along the genome. Peak locations were determined using the MACS algorithm (v1.4.2) with a cutoff of *p* value = 1×10^−7^ (Zhang et al., 2008). ROSE was used to identify enhancers (Whyte et al., 2013). To generate scatterplots of H3K27ac in promoter or distal enhancer regions, the parameters for differential peak calling applied are the following: FPKTM >10 for at least one of the two comparing groups, and FC >1.5.

### Enhancer motif analysis

For H3K27ac ChIPseq, HOMER was used to create tag directories and to call broad peaks. Cisgenome was used to find the distribution of H3K27ac peaks in the genome. We used a cutoff of +/− 3 kb from the TSS to separate enhancers and promoters. *De novo* motif analysis was done using HOMER in the troughs of differentially acetylated enhancers identified from H3K27ac. To compare peak metrics between samples, common intervals are categorized as “Active Regions”, defined by the start coordinate of the most upstream interval and the end coordinate of the most downstream interval. Active regions also include those intervals found in only one sample. Tag density represents DNA fragment per 32-bp bin. The parameters for differential peak calling are the following: <u>F</u>ragments <u>P</u>er <u>K</u>ilobase of DNA per <u>T</u>en <u>M</u>illion mapped reads (FPKTM) >30 for at least one of the groups, and <u>F</u>old <u>C</u>hange (FC) >1.5.

### RNAseq

We isolated total RNA Nucleospin RNA kit (Macherey-Nagel) with DNase I treatment, and followed previously described protocol (Kuo et al., 2014). Directional Poly-A RNAseq libraries were prepared and sequenced as PE42 (42-bp paired-end reads) on Illumina NextSeq 500. For the analysis, 29 to 45 million read-pairs per sample were used. The reads were mapped to the mouse genome mm10 using STAR (v2.5.2b) (Dobin et al., 2013) algorithm with default settings. FeatureCounts from the Subread (v1.5.2) (Liao et al., 2014) package was used to assign concordantly aligned reads pairs to RefSeq genes. A 25-bp minimum as overlapping bases in a fragment was specified for read assignment. Raw read counts were used as input for DESeq2 (v1.14.1) (Love et al., 2014), which was further used to filter out genes with low counts, normalize the counts using library sizes, and perform statistical tests to find significant differential genes. Differential calling results were presented in Supplemental Table S1-6. For statistical analysis, a cutoff of *p* value < 0.05 and FDR < 0.1 was applied, and the resulting genes were used as input for Gene ontology analyses (Ingenuity Pathway Analysis software, Qiagen). Raw and processed data were deposited into the MINSEQE-compliant National Center for Biotechnology Information Gene Expression Omnibus database.

### Fura-2 AM imaging of whole islet secretagogue-stimulate calcium influx

Purified islets were loaded with 2 μM Fura-2-acetoxymethyl ester (AM) for 25 min at 37°C with 5% CO_2_. Islets were washed twice with Kreb’s buffer, and incubated in Kreb’s buffer supplemented with 2 mM glucose for 20 min to reach basal condition before imaging. Then, 14 mM glucose and 45 mM KCl were added sequentially to stimulate calcium flux. Relative intracellular calcium concentrations were measured every 5 sec as a ratio of Fura-2 fluorescence excitation at 340 and 380 nm (F340/F380) both with 510 nm emission, using a Nikon Ti2 microscope equipped with a Photometrics Prime 95B camera, followed by data analysis using Nikon Element software (Dickerson et al., 2018).

### Statistics

Two-tailed Student’s t-test and ANOVA was performed with Prism (GraphPad) for quantitative PCR experiments, glucose tolerance tests, hyperglycemic clamp studies, and secretagogue-stimulated insulin secretion.

### Study approval

The Columbia University Institutional Animal Care and Utilization Committee approved all experiments.

## Acknowledgments

We give special thanks to Mitchell A. Lazar (University of Pennsylvania) and Manashree Damle (Massachusetts General Hospital) for their bioinformatics support and guidance. We also thank members of the Accili laboratory for discussions, and Thomas Kolar, Ana Flete-Castro, and Qiong Xu (Columbia University) for technical support. This work was supported by NIH grants T32DK07328 (to T.K.), K01DK114372 (to T.K.), DK097392 (to D.A.J.), DK115620 (to D.A.J.), DK64819 (to D.A.), DK63608 (to Columbia University Diabetes Research Center), and JPB Foundation (to D.A. and Mitchell A. Lazar).

## Author Contributions

Conceptualization, T.K. and D.A.; Methodology, T.K. and D.A.; Investigation, T.K., W.D., Y.M., P.K.D.; Formal analysis, T.K., W.D., Y.M., P.K.D., D.A.J., and D.A.; Writing-original draft, T.K. and D.A.; Writing-review and editing, T.K., W.D., Y.M., P.K.D., D.A.J., and D.A.; Funding acquisition, T.K., D.A.J., and D.A.

## Supplemental Materials

**Supplemental Figure S1.**
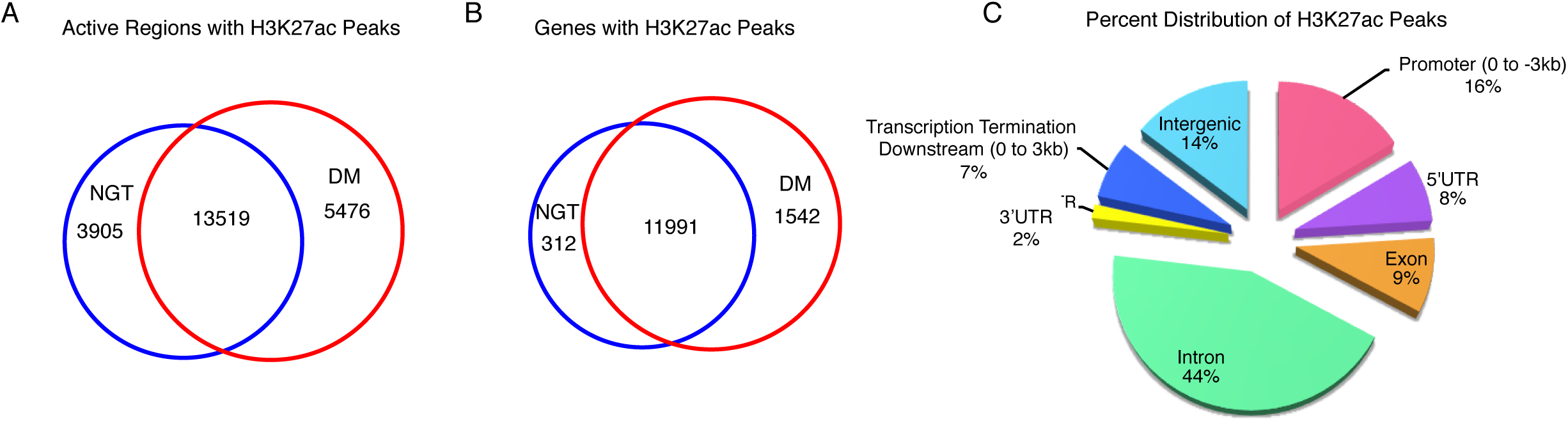
Increased occupancy of H3K27 acetylation in DM β-cells. **(A-B)** Venn diagram showing the genome-wide overlap between NGT and DM for **(A)** H3K27ac active regions and **(B)** genes with H3K27ac peaks. **(C)** A representative pie chart of percent distribution of H3K27ac peaks.

**Supplemental Figure S2.**
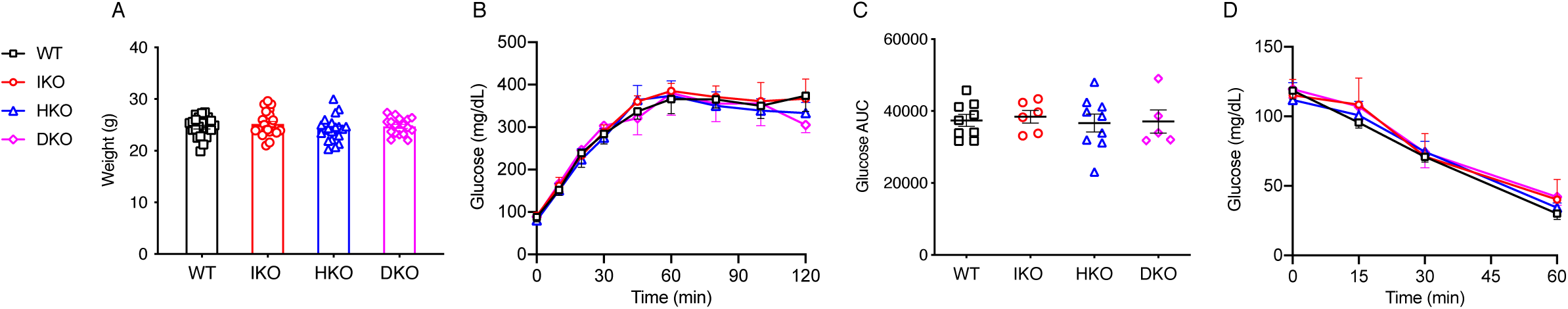
Clamp glucose levels and insulin tolerance tests. **(A)** Average body weight of 6-month old WT (n=11), IKO (n=13), HKO (n=12), and DKO (n=10) mice. **(B-C)** Glucose levels during hyperglycemic clamps. (B) Glucose levels during hyperglycemic clamps in WT (n=9), IKO (n=6), HKO (n=9), and DKO (n=5) mice, and (C) quantification of AUC of B. **(D)** Insulin tolerance test in WT (n=8), IKO (n=5), HKO (n=4), and DKO (n=5) mice.

**Supplemental Figure S3.**
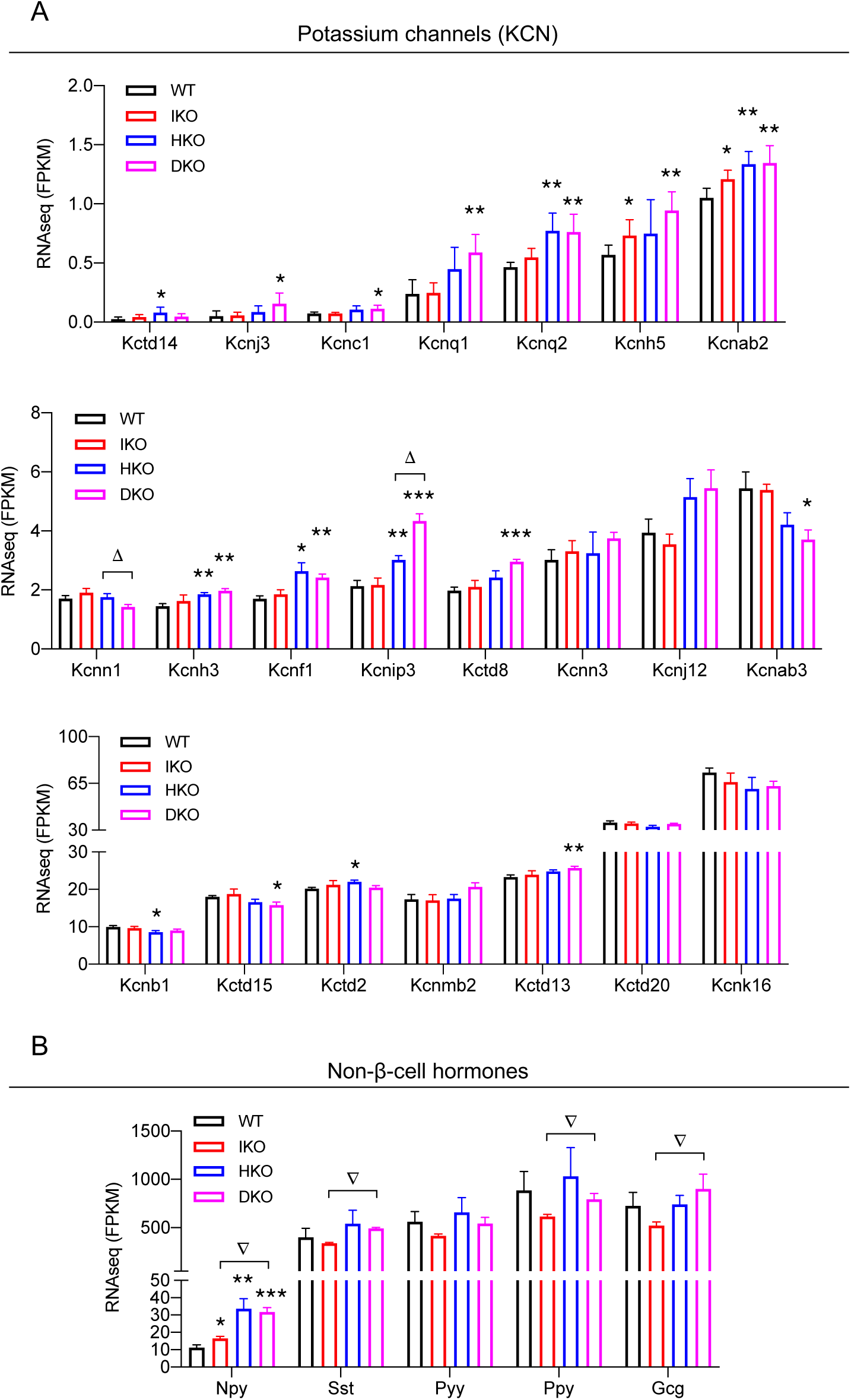
Gene expression profiles for **(A)** potassium channels and **(B)** non-β-cell hormones in WT, IKO, HKO, and DKO β-cells. \**p* < 0.05, \*\**p* < 0.01, \*\*\**p* < 0.001 to WT. Δ *p* < 0.05 for HKO *vs.* DKO. ∇ *p* < 0.05 for IKO *vs.* DKO. n=5 for each genotype.

**Supplemental Figure S4.**
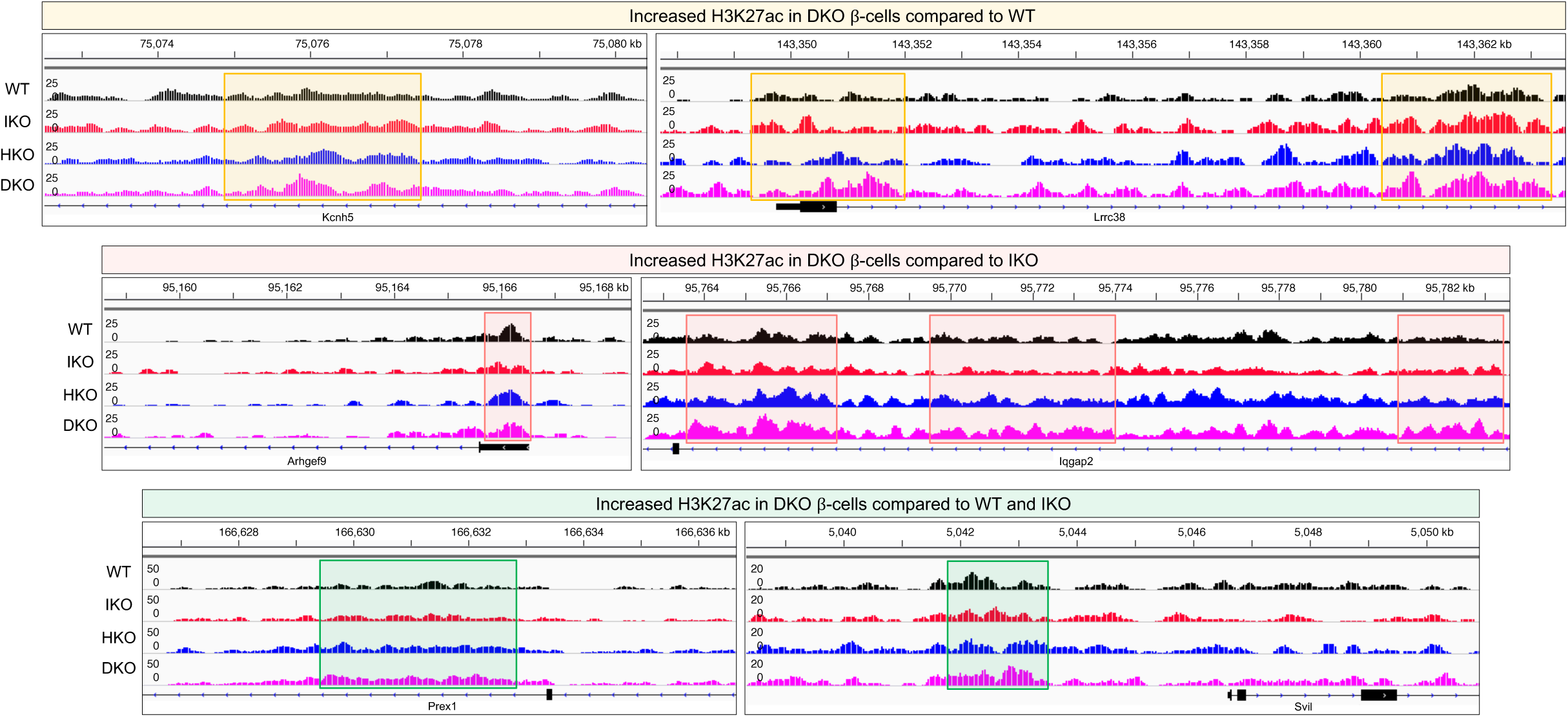
Representative examples of increased H3K27ac levels in DKO β-cells compared to WT, IKO, or both.

## List of materials

### Supplemental Tables

<u>Supplemental Table S1</u>: Differential gene expressions in WT vs. IKO beta cells.

<u>Supplemental Table S2</u>: Differential gene expressions in WT vs. HKO beta cells.

<u>Supplemental Table S3</u>: Differential gene expressions in WT vs. DKO beta cells.

<u>Supplemental Table S4</u>: Differential gene expressions in IKO vs. DKO beta cells.

<u>Supplemental Table S5</u>: Differential gene expressions in HKO vs. DKO beta cells.

<u>Supplemental Table S6</u>: Differential gene expressions in IKO vs. HKO beta cells.

<u>Supplemental Table S7</u>: Concordant RNA and H3K27ac profiles in WT vs. DKO beta cells.

<u>Supplemental Table S8</u>: Concordant RNA and H3K27ac profiles in IKO vs. DKO beta cells.

